# Daisyfield gene drive systems harness repeated genomic elements as a generational clock to limit spread

**DOI:** 10.1101/104877

**Authors:** John Min, Charleston Noble, Devora Najjar, Kevin M. Esvelt

**Author notes:** ***Author‘s Note on Experimental Pre-Registration:*** This manuscript is an example of pre-registration to ensure transparency in experimental gene drive research. It’s intended as a “living document” that begins by sharing key concepts, rationale, and experimental plans for viewing and comment by the community before any experiments begin. As data are gathered and analyses completed, it will be updated with new figures, and eventually will become one or more peer-reviewed publications. All clinical trials now require pre-registration, and the “registered report” model is gaining traction in psychology, but we’re not aware of similar efforts in applied science. This format, which is very much a work in progress, seeks to minimize experimenter effort by making it easy to turn grant proposals into pre-registrations and vice versa while also laying the groundwork for eventual formal publication. Preprint servers offer a way to share the work for external comment while making the document immediately citable in the scientific literature. Openly sharing experimental proposals should accelerate research by allowing scientists to choose whether to collaborate or compete intelligently. For example, if any readers are interested in pursuing these ideas, we would be more than happy to advise, collaborate, or desist in our own efforts as appropriate; we have no desire to duplicate the work of others given the many other urgent projects available. Even if the work is never published in a peer-reviewed journal, the data will remain in place to guide future research along similar lines. It’s worth re-emphasizing that pre-registration preprints are readily transformed into grant proposals and publications alike, so the effort of composition is by no means wasted. If peer evaluation of proposals becomes common, this will not only serve to improve experimental designs and increase safety, but could also increase popular or even financial support of the research. In particular, many small-scale philanthropists are interested in backing promising science, but do not have scientific advisors to assess promising proposals; community evaluations of pre-registrations could potentially lead to funding as well as improve the experimental design. Because gene drive systems could alter the shared environment, we believe that all research in the field should be open to ensure that people have a voice in decisions that could affect them. We hope that our colleagues in gene drive research will join us in sharing their experimental plans; however, we understand that scientists also have moral obligations to their students, whose careers are at risk when outsiders “scoop” their best ideas and publish first. We’re currently in discussions with scientific journals, funders, policymakers, and intellectual property holders concerning ways to change the scientific incentives governing gene drive research so that researchers can freely share their ideas and plans without fear. If the results from open gene drive research are encouraging, we hope that some future descendant of this pre-registration model will spread to the rest of the experimental sciences. **Funding status:** The bulk of this work is currently unsupported. We are grateful for a Burroughs Wellcome Fund “Innovations in Regulatory Science” award to study the evolutionary dynamics of gene drive systems in nematode worms, which will help us evaluate daisyfield once created. Our team has applied to the DARPA Safe Genes program to cover development in nematodes and mosquitoes. **Related projects:** Daisy-chain gene drive.

## Abstract

Methods of altering wild populations are most useful when inherently limited to local geographic areas. Here we describe a novel form of gene drive based on the introduction of multiple copies of an engineered ‘daisy’ sequence into repeated elements of the genome. Each introduced copy encodes guide RNAs that target one or more engineered loci carrying the CRISPR nuclease gene and the desired traits. When organisms encoding a drive system are released into the environment, each generation of mating with wild-type organisms will reduce the average number of the guide RNA elements per ‘daisyfield’ organism by half, serving as a generational clock. The loci encoding the nuclease and payload will exhibit drive only as long as a single copy remains, placing an inherent limit on the extent of spread.

## Popular Summary

Ideally, local communities should be able to change their own environments without imposing those choices on others. Here we describe ways of making a local form of gene drive system for population alteration that can only spread in the presence of “daisy” elements that are initially scattered in many places throughout the genome. When a “daisyfield” organism mates with a wild counterpart, the offspring will typically have only half as many daisy elements as their parent. This creates a stable generational clock that ensures the drive system will eventually stop spreading through the population. Limiting the effects to local populations will enable ethical use by communities who want to solve their own problems without forcing their choices on others.

We haven’t performed any experiments involving daisyfield drive systems yet. Rather, we’re describing what we intend to do, including the safeguards we will use and our assessment of risks, in the hope that others will evaluate our plans and tell us if there’s anything wrong that we missed. We hope that all researchers working on gene drive systems - and other technologies that could impact the shared environment - will similarly pre-register their plans. Sharing plans can reduce needless duplication, accelerate progress, and make the proposed work safer for everyone.

## Introduction

CRISPR-based gene drive systems spread through populations by cutting a target wild-type DNA sequence and causing the cell to copy the engineered drive system and associated genes in its place^1^. However, these ‘global’ drive systems are self-sustaining and should be assumed to spread to every population of the target species in the world, which renders them unsuitable for most applications^2^.

We previously described ‘daisy-chain drive’ systems, which separate the CRISPR components into a series of linked genetic daisy elements such that each element promotes copying of the next^3^. Because the element at one end of the chain is not copied, it is only inherited by half of offspring, whose own offspring have only a 50% chance of inheriting the next element, and so on until all the daisy elements are lost and the ‘payload’ ceases to be copied (Fig. 1). The extent of spread can be programmed by changing the number of daisy elements in the chain and the number of released organisms.

**Figure 1.**
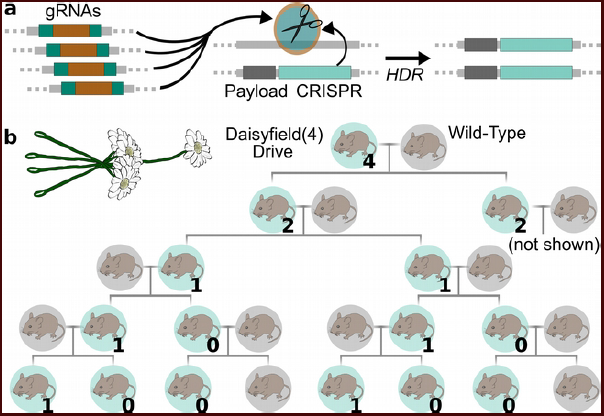
Daisyfield drive systems employ multiple daisy elements encoding the same guide RNAs. (a) A simple four-element daisyfield has four elements that target the wild-type locus harboring the payload and CRISPR nuclease. Cutting and subsequent homology-directed repair (HDR) copies the payload and nuclease. (b) Family tree of the simple 4-element daisyfield assuming each organism mates with wild-type and drive always occurs. The number indicates the copy number of the daisyfield.

One disadvantage of daisy-chain systems is the requirement for numerous CRISPR-based cutting and copying events to occur in every generation. While cutting efficiency typically approaches 100% if multiple guide RNAs are used^4,5^, the homologous recombination rate has varied from 87-99% in the global CRISPR drive systems constructed to date^6–9^. Incorrect repair at multiple loci reduces the overall efficiency of the daisy drive system by generating drive-resistant alleles that prevent inheritance of an intact daisy drive system; such alleles can also segregate away from the payload. Consequently, a form of local CRISPR-based drive that requires only a single cutting and copying step could be useful for many applications, especially if it does not require many separate genome editing events.

## Daisyfield drive systems

Any given gene can be driven by more than one daisy element. Suppose that we add four parallel daisy elements that all drive the same payload element (Fig. 1a). If the elements are unlinked, the next generation will inherit an average of two copies, and will also copy the payload (Fig. 1b). Releasing such an organism is roughly equivalent to releasing four times as many organisms with a standard two-element daisy-chain drive. In general, releasing organisms with 2^N^ copies – a ‘field’ of daisy elements – ensures that all descendants will inherit the payload for (N+1) generations on average, even if each organism exclusively mates with a wild-type counterpart.

The primary advantage of daisyfield drive over daisy-chain drive is that daisyfield requires only a single cut-and-copy event. Because the daisyfield elements are comprised entirely of guide RNAs, the fitness cost to the organism of expressing them should be quite small even when present in large numbers. Only one strong promoter to drive guide RNA expression is required, and there may also be a reduced risk of recombination events capable of creating a global drive system.

Daisyfield is conceptually similar to ‘multi-locus assortment’, in which a desired trait is encoded at multiple sites in the genome of the target organism^10^. Releasing an organism with four copies is theoretically four times as effective as releasing an organism with just one copy, at least as long as any fitness costs due to protein dosage are resolved through the use of feedback. By maintaining the payload at just one locus in all organisms, the daisyfield approach avoids this complication; it is also a true drive system in which the frequency of the payload element will increase in the population over time.

## Combining daisyfield with other drive systems

It’s important to note that daisyfield drive systems are entirely compatible with the daisy-chain drive systems we previously detailed^3^. Conceptually, daisy-chain drive systems are roughly analogous to multi-stage booster rockets (Fig. 2a). Following this analogy, daisyfield drive systems have multiple parallel boosters, half of which run out of fuel and are lost in each generation of mating with wild-type (Fig. 2b). Combining them sacrifices the simplicity of the pure daisyfield drive system, which requires only a single cutting and copying event, but can greatly increase the potency. The most readily constructed version is a daisy-chain drive system for which the normally non-driving element at the end is replaced by a daisyfield (Fig. 2c).

**Figure 2.**
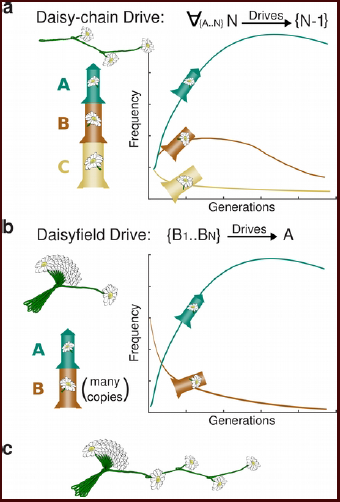
Conceptual analogies of different daisy drive systems. (a) If daisy-chain drives are roughly analogous to multi-stage rockets, then (b) daisyfield drives are tantamount to using multiple parallel boosters, half of which run out of fuel and are lost in each generation of mating to wild-type organisms. (c) The two strategies are most readily combined by using a daisyfield to drive a daisy-chain.

A single daisy-chain element can be also added to drive the daisyfield elements, all of which share a target site. This is only likely to be useful when locating a daisy element on a unique heterogametic sex chromosome, such as the Y chromosome in mammals, to facilitate maintenance of the intact daisyfield drive system in the male line. The major downside of driving the daisyfield elements is that the total number of editing events per generation increases dramatically, which could incur a fitness cost, and that not all daisyfield elements will be successfully copied each generation. Overall, using daisyfield elements to drive a short daisy-chain may be the most potent and therefore economical combination (Fig. 2c).

## Accomplishing efficient multiplex insertion

Multi-locus assortment has never been put into practice, most likely because it suffers from a fundamental engineering problem common to the pre-CRISPR era: too many unlinked elements must be inserted into the genome to be practical.

We believe the solution is to target repeated regions of the genome with CRISPR while supplying a high concentration of donor cassettes with homology to either side of the repeat element (Fig. 3). Repeat targeting has been previously demonstrated in pigs, in which 62 endogenous retroviral elements (ERVs) were removed in just one editing step^11^. If homology-directed repair (HdR) occurs 1/4 as often as non-homologous end-joining (NHEJ) and a viable repair template had been added, 12 copies would be inserted in a single editing event. Most species have numerous candidate repeats. For example, strains of *Mus musculus* has ~10-40 copies of murine leukemia virus^12^, ~85 copies of VL30^13^, and ~200-300 copies of EtN^14^.

**Figure 3.**
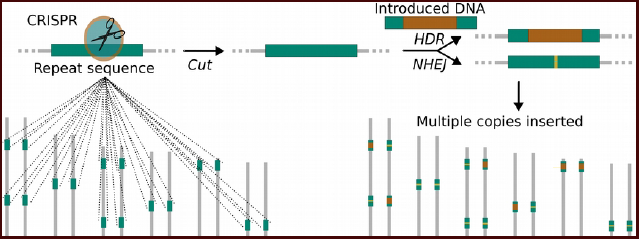
Targeting a repeated element with CRISPR could insert many copies of an introduced element in a single step through homology-directed repair (HDR). Other copies would acquire mutations through non-homologous end-joining that block further cutting by that guide RNA.

## Constructing high-copy-number daisyfield systems

Building highly potent daisyfield drive systems will require us to generate stable strains with dozens or hundreds of insertions. This poses a formidable challenge given typically low HDR rates outside the germline and early embryo. Each organism generated using the repeat insertion strategy is likely to have copies of the donor sequence inserted at a different subset of the target repeats, with most or all other copies of the target sequence eliminated via NHEJ. Maximizing the number of engineered repeats will require a means of converting NHEJ-generated mutant repeats to the desired sequence.

One of the best ways to repeatedly convert unwanted sequences to desirable ones is to use a daisy drive. For example, one might stably introduce a DNA cassette that encodes a fluorescent marker, a germline-expressed CRISPR nuclease gene, and one of several different guide RNAs targeting the wild-type repeat sequence (Fig. 4a). Co-delivering this cassette along with the DNA sequence to be inserted into the repeat will produce organisms carrying the cassette, repeats with the desired sequence, and repeats with NHEJ-generated mutations. Crossing organisms that carry different guide RNAs in the stable cassette will result in heterozygotes in which the nuclease will cut all NHEJ mutants in the germline, affording another chance to convert NHEJ alleles into repeats when HDR rates are at their highest (Fig. 4b). The resulting progeny can be crossed with equivalents carrying still different guide RNAs to provide more opportunities for conversion (Fig. 4c), while outcrossing to wild-type can help maintain fitness while driving most daisyfield elements. The dominant visible marker provides a way to remove the cassette harboring the maintenance nuclease before deployment.

To maximize the total number of knock-ins, ssDNA oligonucleotides might be used to insert recombinase recognition sites into the repeats with higher efficiency than longer dsDNA templates can offer. This approach would enable guide RNA cassettes for any daisyfield strain, or any payload if using multi-locus assortment, to be subsequently inserted via recombinase-mediated cassette exchange (Fig. 4d).

**Figure 4.**
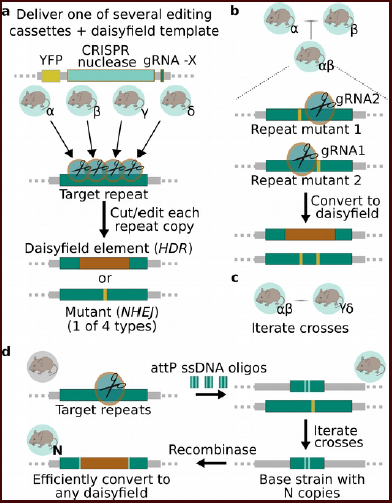
A strategy to generate high-copy-number daisyfield strains. a) Stably inserting different versions of a CRISPR-encoding cassette, each with a different guide RNA targeting the repeat sequence, into the genome when introducing the daisyfield elements will create parallel strains with different repeat insertion patterns. b) Crossing two lines will convert mutant alleles generated by non-homologous end-joining (NHEJ) to daisyfield elements by homology-directed repair (HDR). c) Iterating this process and out-crossing to wild-type to increase fitness while screening for animals with the most copies can maximize the number of daisyfield elements. d) Initial insertion efficiency can be maximized and adaptability preserved by using this strategy ssDNA oligonucleotides encoding recombinase sites into the repeats, then delivering the desired daisy elements via recombinase-mediated cassette exchange.

## Discussion

Scattering daisy elements throughout the genome can generate a ‘daisyfield’ drive systems that will spread through local populations, but is limited by a satisfyingly simple generational clock: the average number of daisy elements will be halved after each generation of mating with wild-type organisms. Once none are left, the payload will be inherited normally.

Satisfyingly, daisyfield drive systems might be generated by replacing ancient selfish genetic elements scattered throughout the genome, thereby replacing a selfish genetic network with an altruistic one. Crossing strains using different guide RNAs for insertion would permit the construction of highly fit strains with many copies yet otherwise wild-type genetics. These could be subsequently converted into any desired daisyfield drive system through highly efficient recombinase-mediated cassette exchange.

Which local CRISPR-based drive system is superior: daisy-chain drive, daisyfield drive, or a combination of both? The answer likely depends on both the target organism and the application. For example, organisms that are particularly amenable to inserting many copies at once or prone to recombination may favor the daisyfield approach. In contrast, applications involving population suppression through sex-specific infertility may want to place the initially non-driving element on the chromosome of the opposite sex for easier maintenance of a daisy-chain, while drive systems intended to affect very large populations may need a combination to maximize potency.

Mathematical models of different types of daisy drive systems will shed light on their comparative benefits, particularly when combined with experimental studies of evolutionary dynamics and stability in large populations. We consequently plan to build and study all types of daisy drive systems in nematode worms. In keeping with our support for the pre-registration of experimental plans involving gene drive, we have detailed our intended series of experiments and safeguards below. Our hope is that the community will embrace pre-registration and help improve our designs and safety measures through feedback. We will update this manuscript with more detailed experimental plans as they are designed, as well as the results of mathematical modeling and analyses of experimental data.

### Experimental Pre-Registration

<u>Nematodes:</u> *C. elegans* has a fully sequenced and assembled genome and is easy to engineer via microinjection, but undergoes transgene silencing in the germline. We will accordingly test multicopy insertion in this worm, but may not be able to test the full daisyfield system unless nuclease licensing efforts succeed. Purified Cas9 complexed with guide RNAs and a high concentration of ssDNA oligonucleotides will be delivered via computer-assisted microinjection of live worms using our custom-built apparatus. We will target the Cele1 (~1000 copies) element as well as repeats with ~100 and ~20 copies, to be determined by repeat analysis^15^. The copy number of the insert will be measured by qPCR. *C. brenneri* worms are not thought to undergo germline silencing, but their genome is not fully assembled into chromosomes, which makes evaluating repeat locations more difficult. They may also have notably different transposable elements. We will identify repeats with equivalent copy numbers using repeat analysis software, check for genome location to ensure no element is in close proximity to the payload.

Payload elements will be ribosomal genes with the last exon recoded and the _3_’UTR swapped for that of a different ribosomal gene. The nuclease will be a copy of the AsCpf1 or SpCas9 gene driven by the mex-5 promoter with the *gld-1* _3_’UTR for germline-specific expression^16^. In most cases, the nuclease will be present in the strain background to avoid the risk of recombination that might generate a global drive system. When the goal is to study whether this occurs, the nuclease will be included as part of the payload element, but the drive system will target a synthetic site in the recipient strain. If necessary, we will undertake germline licensing via fusion to exons of already licensed genes such as oma-1^17^ and fbf-1 or by PATC insertion into introns^18^. In all cases, the payload will encode a fluorescence marker gene, and typically eliminate a different marker gene in the recipient strain.

For daisy-chain drive systems, we will insert daisy elements immediately downstream of ribosomal genes in neutral sites, or downstream of recoded ribosomal elements. All daisy elements will drive the payload as well as the next element in the daisy-chain to maintain genetic linkage between each element and the payload.

Worms will be grown with synchronized generations in large flask populations or in wells of culture plates with liquid transferred between plates for controlled gene flow between subpopulations. To examine daisy drive stability, a group of serially linked large populations will be seeded at one end with daisy drive worms and the abundance of each strain monitored over time via plate reader or plating and automated quantification.

<u>Mice:</u> We intend to insert attP sequences into MLV, VL30, and EtN sequences by embryo injection of purified Cas9 and guide RNAs along with ssODN templates. Knock-in rates are substantially lower in the embryo than in the germline, but it is difficult to generate large numbers of DNA templates for repair in vivo. Accordingly, we will rely on delivery by microinjection as soon after fertilization as is feasible, when homology-directed repair rates are relatively high.

<u>Safeguards:</u> Daisyfield is itself a form of molecular confinement; generating a global drive system from a daisyfield system would require a recombination event that moved guide RNAs from a repeat sequence adjacent to the nuclease. We will ensure that the two sequences do not share more than 12 base pairs of homology to prevent homologous recombination, which typically requires 18 or more base pairs, and will try to ensure that the nuclease gene is located at least 100kb away from the nearest repeat sequence targeted for daisyfield insertion (this may be difficult given the *C. brenneri* assembly). Whenever we are not testing the stability of a daisy drive system, the nuclease will be present in a locus that is not targeted by any guide RNAs and instead supplied by the recipient strain of worms. Our nematode studies will additionally employ ecological containment, as *C. elegans* are not found in New England and *C. brenneri* is an exclusively tropical species^19–21^. In mice, we will employ barrier containment: MIT animal facility strictly controls access to the animal facilities, with all animals kept in cages within secured rooms constructed to eliminate any chance of rodent escape.

